# Towards an autologous iPSC-derived patient-on-a-chip

**DOI:** 10.1101/376970

**Authors:** Anja Patricia Ramme, Leopold Koenig, Tobias Hasenberg, Christine Schwenk, Corinna Magauer, Daniel Faust, Alexandra K. Lorenz, Anna Krebs, Christopher Drewell, Kerstin Schirrmann, Alexandra Vladetic, Grace-Chiaen Lin, Stephan Pabinger, Winfried Neuhaus, Frederic Bois, Roland Lauster, Uwe Marx, Eva-Maria Dehne

**Affiliations:** TissUse GmbH Oudenarder Str. 16 13347 Berlin Deutschland; AIT Austrian Institute of Technology GmbH Giefinggasse 4 1210 Vienna Austria; The University of Manchester Physics of Fluids and Soft Matter Group Oxford Road Manchester M13 9PL UK; INERIS, METO unit Parc ALATA BP2 60550 Verneuil en Halatte France; Technische Universität Berlin Medizinische Biotechnologie Gustav-Meyer-Allee 25 13355 Berlin Deutschland

## Abstract

Microphysiological systems are fundamental for progressing towards a global paradigm shift in drug development through the generation of patient-on-a-chip models. An increasing number of single- and multi-organ systems have been adopted by the pharmaceutical and cosmetic industries for predictive substance testing. These models run on heterogeneous tissues and cell types from different donors. However, a patient is an individual. Therefore, patient-on-a-chip systems need to be built from tissues from one autologous source. Individual on-chip organ differentiation from a single induced pluripotent stem cell source could provide a solution to this challenge.

We designed a four-organ chip based on human physiology. It enables the interconnection of miniaturized human intestine, liver, brain and kidney equivalents. All four organ models were predifferentiated from induced pluripotent stem cells from the same healthy donor and integrated into the microphysiological system. The cross talk led to further differentiation over a 14-day cultivation period under pulsatile blood flow conditions in one common medium deprived of growth factors. This model platform will pave the way for disease induction and subsequent drug testing.

## Introduction

The current drug development cycle is inefficient due to the high failure rate of preclinical drug safety and efficacy testing, as well as inadequate translation from preclinical animal models to patients. However, the demand for new effective drugs is increasing, for example, because of lifestyle, environmental changes or rising life expectancy. The latter has led to a greater prevalence of chronic and age-associated health disorders, such as diabetes, rheumatic arthritis, Alzheimer and Parkinson’s disease. Consequently, no cure for these diseases has been discovered despite decades of drug research. Personalized multi-organ microphysiological system (MPS) models, based on patient-derived induced pluripotent stem cells (iPSC) represent a promising approach to address this^1, 2^.

The MPSs are capable of emulating human biology *in vitro* at the smallest biologically acceptable scale. The dynamic fluid flow is adjusted to enable physiological nutrition and oxygen supply of the tissues mimicking organ function with a minimum use of human cells and tissues^3^. The technology of multiorgan MPSs can mimic complex biological processes involving organ-organ interaction, system homeostasis and pharmacokinetics. These systems will enable quicker, more accurate, cost-effective and clinically relevant testing of drugs^3, 4^.

A lot of stem cell-based approaches for cell differentiation have emerged in recent years and several reviews regarding the different perspectives on combining stem cells and MPS have been published^1, 2, 5–8^: (i) functional differentiation of iPSCs into cardiomyocytes or hepatocytes on a chip^9^; (ii) differentiation and cultivation of three-dimensional (3D) brain organoids in a perfusable organ-on-a-chip system^10^; and (iii) mesenchymal stem cell cultures and differentiation in microfluidic devices^5^. However, to the best of our knowledge, there is no multi-organ MPS co-culturing solely iPSC-derived organ equivalents in a common media circuit. A model that recapitulates the effect of a disease on iPSC-derived organoids from a healthy donor could be a promising assay platform. Moreover, there is great potential to generate tailored disease models by using iPSC-derived organoids from a disease donor or genome editing on healthy iPSCs.

Conventional iPSC-derived cultures require highly specialized culture conditions. Each cell type, for example, demands its own set of adjusted growth factors. The co-culture of several organ equivalents is not straightforwardly possible using routine culture methods. To address this issue, we adapted our recently published microphysiological multi-organ-chip (MOC) platform to host the organ equivalents desired^11–13^. It has been shown previously that the MOC can co-culture tissue models of various origins, based on cell lines or primary cells, for up to 28 days under homeostasis^11, 12^. We assumed that predifferentiated organ models for the intestine, liver, brain and kidney of iPSC origin could maintain their phenotype similarly under the culture conditions of the microfluidic device.

We here describe for the first time the co-culture of four iPSC-derived organ equivalents from one single donor on the MOC platform. We developed a comprehensive ADME (absorption, distribution, metabolism and excretion) MPS with the area of a standard microscopic slide for the long-term culture of human iPSC-derived organ equivalents (termed “ADME-MOC”). The organ models introduced not only maintained their phenotype during the 14 days of co-culture in a common, growth factor-deprived medium, but also further differentiated physiologically along the initiated path.

## Materials and methods

### ADME-MOC fabrication

Standard soft lithography and replica molding of polydimethylsiloxane (PDMS; Sylgard 184, Dow Corning) were applied for the fabrication of the chip. Briefly, two master molds were fabricated by aluminum milling – one mold for each circuit. An adapter plate (containing access holes) was treated with a silicon rubber additive (1200 OS Primer, Dow Corning) and clamped to the master mold for the excretory circuit. Liquid PDMS (10:1 v/v ratio of PDMS to curing agent) was injected into this casting station. Screws were used to generate the PDMS-free culture compartments and the PDMS membranes constituting the two on-chip micropumps. The setup was incubated at 80 °C for at least 60 min. The PDMS slice bonds fluid-tight to the adapter plate. A second PDMS slice of the bottom blood circuit was prepared without an adapter plate and without through holes. After curing, the PDMS slices were treated with low-pressure ambient air plasma (Diener GmbH, Pico) and immersed in a 2 % (3-Glycidyloxypropyl) trimethoxysilane aqueous solution (440167-100ML Sigma-Aldrich) for 20 min at room temperature. In parallel, a precut cell culture ready-to-use polycarbonate membrane (pore size: 1.0 μm, pore density: 2*10^6^ cm^-2^, thickness: 24.0 μm, it4ip S.A.) was also treated with plasma and immersed in 1 % (3-Aminopropyl) triethoxysilane aqueous solution (A3648 Sigma-Aldrich) for 20 min at room temperature. After drying, the membrane was placed on the space provided on the bottom PDMS slice exactly at the position of the glomerulus and tubules compartments. Both PDMS slices were aligned. Critical PDMS membranes were drawn upwards using a vacuum of less than −20 kPa to prohibit their bonding. The chip was then incubated for at least 24 h at 80 °C, which realized a fluid-tight bonding of the two PDMS slices. The chips were rinsed with ethanol, phosphate buffered saline (PBS) and, subsequently, cultivation medium. Thereafter, the chip was kept at 37 °C in a humid incubator until the start of the experiment.

### Characterization of fluid dynamics

The two integrated peristaltic micropumps realized the flow in both microfluidic circuits. Their working principle is explained elsewhere^14^. The deflection of the pumps’ membranes was controlled by an external unit (TissUse GmbH) that managed the frequency of deflection, pressure and vacuum. Their interplay defined the flow rate within the two circuits. Micro particle image velocimetry (μPIV) was applied to determine the flow rate for the set process parameters. Red blood cells were utilized to visualize the flow. Blood was obtained from human donors by venipuncture. After centrifugation at 3.0 *g* for 10 min, the plasma was discarded and the red blood cells were re-suspended in PBS. The hematocrit was set to 2.5 % for the image acquisition. The μPIV analyses were carried out in straight channels. Throughout the measurements, the chip was equipped with the same cell culture inserts and volumes as in the corresponding biological experiments. The flow was recorded with a high-speed CMOS camera (HXC40, Baumer Optronic) coupled to an inverted microscope (Axiovert, Zeiss). Magnification was set to either 2.5 or 5x, which resulted in a resolution of 0.23 and 0.46 px/μm, respectively. The acquisition rate’s upper limit was 2,583 fps. The displacement of the red blood cells in recordings of up to 7.7 s was computed by the open source toolbox “PIVlab”^15, 16^. Only velocity vectors in the central portion of the channels were regarded to determine the flow rate of the laminar plug flow. Details of the underlying equations have been discussed elsewhere^14^. The flow rates were averaged from triplicate measurements. The standard deviation is given.

### iPSC generation and cultivation

Cell culture plates and components were purchased from Corning U.S. and cultures were incubated at 37 °C and 5 % CO_2_, unless otherwise stated. The human iPSC line StemUse101 (TissUse GmbH) was derived from peripheral blood mononuclear cells from a 52-year-old male donor. Human blood samples were donated with informed consent and ethics approval (Ethic Committee Berlin Chamber of Physicians, Germany) in compliance with the relevant laws. Reprogramming was performed by Phenocell SAS with an episomal vector (Epi5 Kit, Thermo Fisher Scientific A15960). The iPSCs were maintained in feeder-free conditions in StemMACS iPS-Brew XF (Miltenyi) on growth factor-reduced (GFR) Matrigel^®^ (1:100 diluted in KO/DMEM F12; Thermo Fisher Scientific) on dishes treated for cell culture. The iPSCs were passaged every five to seven days using Accutase^®^, 4,000 – 6,000 cells/cm^2^ were seeded and medium was added with 10 μM Rock Inhibitor Y-27632 (Cayman). StemMACS iPS-Brew XF medium without Rock Inhibitor was renewed after 48 h, following a daily medium exchange. The iPSC differentiations were performed with cells in passage 18 to 25.

### Differentiation into stromal cells

Stromal cell differentiation was carried out with modifications from Zou et al.^17^. The iPSCs were grown on GFR Matrigel until 80 % confluent. The medium was changed stepwise to stromal cell medium (50 % DMEM HG with 50 % Ham’s F-12, 10 % FCS, 15 mM HEPES and 1 % penicillin-streptomycin: P/S) with one transitional day of 50:50 stromal cell medium with StemMACS iPS-Brew XF. The medium was changed daily during the first four days and, subsequently, renewed every two or three days. After three weeks, cells were split 1:3 with trypsin on 0.1 % gelatin (Sigma)-coated cell culture dishes; 10 μM Rock Inhibitor Y-27632 was added to the medium only for the split. Cells were split every two weeks or when confluent. The iPSC-derived stromal cells in passage four were used for the intestine and liver models.

### Differentiation into hepatocytes and liver spheroids formation

Hepatocyte cell differentiation was carried out with modifications from Szkolnicka et al.^18^. The iPSCs were differentiated into definitive endoderm (DE) with STEMdiff™ Definitive Endoderm Kit (TeSR™-E8™ Optimized) (StemCell), according to the manufacturer’s instructions with minor modifications. The iPSCs were split with Accutase and seeded with 33,000 cells/cm^2^ cells on GFR Matrigel (1:100 diluted in KO/DMEM F12) in StemMACS iPS-Brew XF supplemented with 10 μM Rock Inhibitor and STEMdiff™ Definitive Endoderm TeSR™-E8™ Supplement; 1:20. Supplement A and B treatment was performed two days after seeding. After 24 h, only supplement B was added for three days, as described in the instructions. The cell layer was then treated for five days with 1 % dimethyl sulfoxide in KO/DMEM (Thermo Fisher Scientific), 20 % KnockOut™ Serum Replacement (KSR) (Thermo Fisher Scientific), 1 mM Glutamine, 1 % nonessential amino acids, 0.1 mM 2-mercaptoethanol (Thermo Fisher Scientific) and 1 % P/S. Due to pronounced cell death during the first two days of the dimethyl sulfoxide treatment, the cells were gently washed with PBS before the medium was replaced. Afterwards, the cell layer was treated for an additional five days with HZM medium (HepatoZYME-SFM (Thermo Fisher Scientific) medium with 1 mM nonessential amino acids, 2 mM L-glutamine, 2 % KSR, 10 ng/mL human HGF (Miltenyi), 10 ng/mL human FGF-4 (Peprotech), 10 ng/mL human oncostatin M (Miltenyi), 0.1 μM dexamethason (EHRENSTORFER GMBH) and 1 % P/S). Gentle medium exchange was performed every other day. Liver spheroids were formed combining iPSC-derived hepatocytes and iPSC-derived stromal cells using Corning^®^ 384-well Spheroid Microplate. The iPSC-derived hepatocytes and iPSC-derived stromal cells were detached with trypsin, centrifuged and diluted in HZM medium (described previously) supplemented with 10 μM Rock Inhibitor and 5 % Bovine Serum Albumin Fraction V, fatty acid free (BSA-FAF) (SERVA). A 50 μL cell suspension containing 4.8 × 10^4^ hepatocytes and 0.2 × 10^4^ stromal cells was pipetted into each access hole with a 96-well pipette (Platemaster^®^, Gilson). After two days of culture on an orbital shaker (Corning), the spheroids were transferred with wide-bore tips to ultra-low attachment 24-well plates. Twenty spheroids were collected together to form a single liver equivalent in the respective culture compartment of the ADME-MOC.

### Differentiation into intestinal organoids and cell culture insert^®^ seeding

Intestinal organoid differentiation was carried out with modifications from Kauffman et al.^19^ and McCracken et al.^4^. The iPSCs were differentiated into DE with STEMdiff™ Definitive Endoderm Kit (TeSR™-E8™ Optimized), as described above. Following the DE stage, the cell layer was treated with hindgut medium (Advanced DMEM/F-12; Thermo Fisher Scientific) with 50 ng/mL KGF (Peprotech), 0. 1 % BSA-FAF, 15 mM HEPES, 1x B27-Supplement (Thermo Fisher Scientific) and 1 % P/S for four days. The hindgut medium was added with 50 ng/mL KGF, 2 % BSA-FAF, 15 mM HEPES, 1x B27-Supplement, 1 % P/S, 1x insulin-transferrin-selenium (ITS), 200 mg/L, ethanolamin (Sigma) and 2 μM retinoic acid (Alfa Aesar) for the following three days. Subsequently, the 3D cell layer was scraped off with a pipette tip and small pieces were transferred with a cooled wide-bore pipette tip into cool 100 % Matrigel (354234). The suspension was mixed gently without producing bubbles and 50 μL Matrigel cell suspension was transferred immediately with a cooled wide-bore pipette tip into the middle of a 37 °C pre-warmed 24-well plate. The Matrigel was solidified in the incubator for 10 min, then 500 μL intestinal medium (Advanced DMEM/F-12 with 2 mM L-glutamine, 15 mM HEPES, 1x B27-Supplement, 1 % P/S, 500 ng/mL R-Spondin-1 (Peprotech), 100 ng/mL Noggin (Peprotech) and 100 ng/mL EGF (StemCell) was added. Intestinal organoids were passaged every two weeks by resuspending the organoids 10x with a 200 μL pipette in PBS, centrifuging at 300 *g* at 4 °C for 3 min and transferring the organoids in Matrigel, as described above. Expansion of organoids was increased stepwise from 1:2 to 1:20. The intestinal medium was changed every two to three days. For the cell culture insert (Millicell PCF 0.4 μm pores PIHP01250) intestinal model, 0.6 x 10^6^ iPSC-derived stromal cells per cell culture insert were seeded in stromal cell medium. After two days, the cell culture inserts were washed gently with PBS, 50 μL cooled Matrigel was added and the intestinal organoids in passage six were seeded on top of the stromal cells in 70 μL Matrigel-medium (50/50) suspension. One Matrigel organoid 24-well droplet was used for two cell culture inserts. After the Matrigel solidified, 100 μL intestinal medium was added into the cell culture insert and 500 μL underneath. Intestinal medium was changed every two to three days. The intestinal models on cell culture inserts were cultivated for two weeks before transferring them into the ADME-MOC.

### Differentiation into renal cells

Renal cell differentiation was carried out with modifications from Morizane and Bonventre^21^. The iPSCs were grown until 50 % confluent. Subsequently, the cells were washed gently with PBS and the medium was replaced by advanced RPMI 1640 (Thermo Fisher Scientific) with 2 mM L-glutamine, 5 μg/mL Gentamycin Sulfate and 0.25 μg/mL Amphotericin B (renal basal medium), and 8 μM CHIR99021 (LC Labs) was added to the medium during the first four days. Medium was replaced every other day. Afterwards, the renal basal medium was added with 10 ng/mL Activin A (Peprotech) for three days without medium exchange. The cells were then washed with PBS and 10 ng/mL FGF-9 (Peprotech) was added to the renal basal medium. Additionally, two days later, 3 μM CHIR99021 with 10 ng/mL FGF-9 induction was carried out. Subsequently, the cells were detached using Accutase and about 10^6^ cells were injected gently into the excretory circuit of each ADME-MOC. The excretory circuit was coated previously with 1 % GFR Matrigel and filled with renal basal medium supplemented with 10 ng/mL FGF-9. Two days later, 50 % renal basal medium supplemented with 10 ng/mL FGF-9 was renewed in the chip. Renal cells were injected six days before the start of the ADME-MOC co-culture.

### Differentiation into cortical neurospheres and Transwell^®^ seeding

Differentiation of iPSCs into cortical progenitor cell spheroids, referred to as neurospheres, was performed in a DASbox^®^ Mini Bioreactor System (Eppendorf). The culture protocol was adapted from Rigamonti et al.^22^. Initial spheroid formation of iPSCs in the DASbox bioreactor was performed as described by Abecasis et al.^23^. The iPSCs were inoculated as a single cell suspension with a concentration of 2.5 x 10^5^ cells/mL in 125 mL StemMACS iPS-Brew XF supplemented with 10 μM Rock Inhibitor Y-27632 in a 250-mL fermentation vessel (DASbox Mini Bioreactor System). The vessel was equipped with a trapezoid-shaped impeller, a submerged tube for media withdrawal, a non-submerged tube for media feeding, and a dissolved oxygen and pH sensor. Sensors and pumps were calibrated according to the manufacturer’s protocols. The submerged sampling tube was equipped with a 10 μm porous filter to allow the washout of single cells while retaining the iPSC spheroids in the vessel. The vessel was kept at a temperature of 37 °C and aerated with 3 sl/hour (5 % CO_2_, variable O_2_) and a stirrer speed of 80 rpm. The aeration with oxygen was adjusted automatically by DASware control software (Eppendorf) to achieve a stable oxygen saturation at 19 % dissolved oxygen. No medium was exchanged for the first 24 h after inoculation. Starting at day two, medium was exchanged via the tubing system at a rate of 120 mL/day at day two, 60 mL/day at day three, 90 mL/day at day four and 120 mL/day at day five. After five days of cultivation, the process was stopped temporarily, and the spheroids were used for cortical differentiation. Agitation was stopped and the spheroids were allowed to settle at the bottom of the vessel. The culture medium was removed completely and the spheroids suspended in a defined volume of fresh iPSC culture medium. The suspension was mixed well and a sample was taken for cell counting with an automated cell counter (Nucleocounter NC-200, Chemometec). Cells were reintroduced as spheroids into the DASbox bioreactor system with 7.5 x 10^5^ cells/mL in 100 mL StemMACS iPS-Brew XF. The system was operated with the same process parameters, as described for the spheroid formation. After 24 h (day two of differentiation), the medium was completely exchanged manually with KSR medium (KO-DMEM (Thermo Fisher Scientific) with 15 % KSR, 1 % nonessential amino acids, 2 mM L-Glutamine, 1 % P/S and 50 μM 2-mercaptoethanol). The medium was constantly exchanged at a rate of 2.08 mL/h from day two on. Only KSR medium was fed until day 4. From then on to day 10, the KSR medium was gradually diluted with neural induction medium (DMEM/F12 (Thermo Fisher Scientific) with 1 % N2 (Thermo Fisher Scientific), 2 % B27 without vitamin A (Thermo Fisher Scientific), 1 % nonessential amino acids, 2 mM L-Glutamine and 1 % P/S. Only neural induction medium was fed from day 10 to day 30. The cell culture medium was supplemented with 10 μM SB431542 (Miltenyi) and 1 μM LDN193189 (Sellekchem) from day one to day seven. From day two to four, medium was furthermore supplemented with 2 μM XAV939 (Cayman). After 30 days, neurospheres were collected and cell debris was removed with a 37 μm reversible strainer (Stemcell). The spheroids were resuspended in a defined volume of neural cultivation medium (Neurobasal (Thermo Fisher Scientific) with 1 % N2, 2% B27 without vitamin A, 1% nonessential amino acids, 2 mM L-glutamine and 1 % P/S. The suspension was mixed well and a sample was taken for cell counting. The neurospheres were further cultivated in neural cultivation medium in an Erlenmeyer flask on an orbital shaker. The next day, the neurospheres were transferred into a 96-well Transwell^®^ system with a 1-μm pore polyester membrane (Corning). A total of 2 x 10^6^ cells was transferred into each Transwell. Spheroids were cultured for 24 h in neural cultivation medium before individual Transwells were transferred into the microfluidic chip system.

### Chip-based co-cultures

The excretory circuits of each chip were loaded with renal cells six days prior to the start of the coculture experiment, as described above. All preloaded chips were equipped with the intestine cell culture insert, liver equivalents and neuronal Transwell model to start the co-culture experiment. A pumping frequency of 0.5 Hz in the surrogate blood circuit and 0.4 Hz in the excretory circuit were chosen to ensure pulsatile fluid flow. The pressure was set to 450 mbar and the vacuum was set to −350 mbar for both pumps.

The following medium was used in the ADME-MOC: Williams E without phenol red with/without glucose (PAN) with the addition of 1 mM nonessential amino acids, 2 mM L-glutamine, with/without 5 % human AB serum, with/without 1x ITS, 20 μg/mL gentamycin sulfate and 1 μg/mL amphotericin B. No human AB serum or ITS was added to the medium in the excretory circuit. A glucose feeding solution was prepared by dilution of a glucose-rich Nutriflex^®^ peri solution (B. Braun) to a glucose concentration of 20 g/L. The pH of the glucose feeding solution was adjusted to 7.2. Initially, the chip contained the following media: 225 μl intestinal medium on the apical side of the intestine model mixed with 50 μl glucose feeding solution, 1.37 ml ADME-MOC medium with glucose, human AB serum and ITS in the surrogate blood circuit, 75 μL neural cultivation medium on the apical side of the brain equivalent and 575 μL ADME-MOC medium with glucose but without human AB serum and ITS in the excretory circuit. Samples were taken from both medium reservoirs (100 μL) and from the apical side of the intestine equivalent (50 μL) daily. The medium removed was replaced with equal volumes of glucose-free medium for the surrogate blood medium reservoir and glucose-, serum- and ITS-free medium for the excretory medium reservoir. At the intestinal compartment, 50 μL glucose feeding solution was applied. In this way, 1.05 mg of glucose was fed daily to the system only through the intestine equivalent. Sets of chips (seven replicates) were cultivated for 7 and 14 days. Cellular samples were taken for immunohistological examinations, qPCR and RNA sequencing at the endpoints. Supernatants were analyzed for their glucose and lactate dehydrogenase (LDH) content.

Tissue viability was monitored daily by the measurement of LDH released in the supernatants of the three media pools: intestinal lumen pool, surrogate blood circuit pool (medium reservoir 1) and excretory circuit pool (medium reservoir 2 (Figure 1 A). The LDH activity of the medium was measured using the Cytotoxicity Detection KitPLUS (Roche), according to the manufacturer’s instructions with minor modifications. In brief, an amount of 12.5 μL of reagent was used and 12.5 μL of the sample was added for each measurement. A standard curve based on the LDH Positive Control (L-LDH Standard stock solution from rabbit muscle, #10127230001, Sigma) was prepared. Samples were incubated in 384-well microtiter plates for 20 min on an orbital shaker. Absorbance readings at 490 nm with 680 nm background correction were performed in a microplate reader (BMG Labtech ClarioStar). If necessary, samples were diluted with 1 % BSA in PBS.

**Figure 1:**
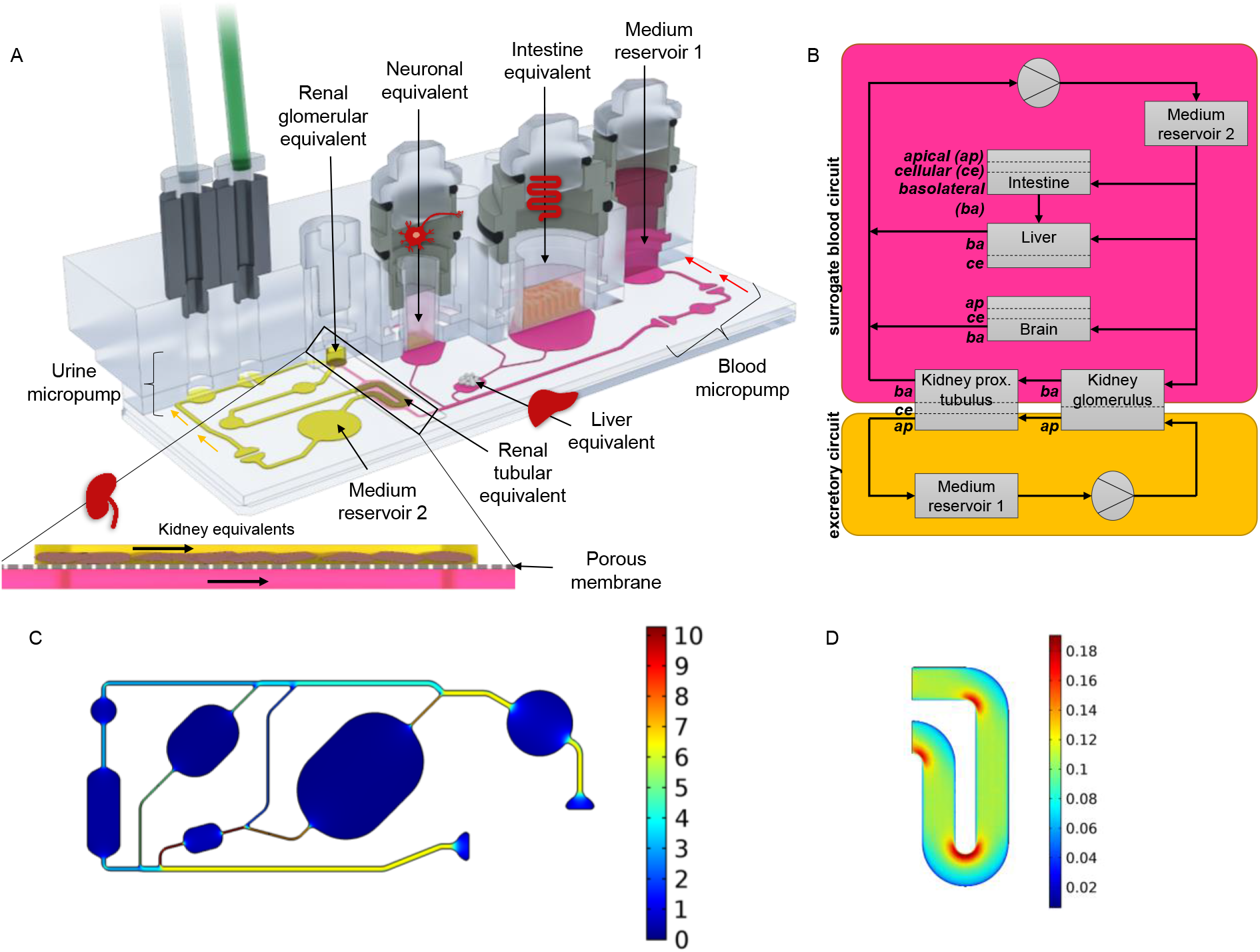
A: The microfluidic ADME-Multi-Organ-Chip (ADME-MOC) at a glance (A), pharmacokinetic/pharmacodynamic (PK/PD) modelling of the ADME-MOC (B). Pink: surrogate blood circuit, yellow: excretory circuit. Calculated velocity distribution (mm/s) at a flow rate of 16.9 μl/min in the surrogate blood circuit (C). Distribution of the wall shear stress (Pa) in the tubular compartment of the excretory circuit at a mean shear stress of 0.1 Pa (D).

The metabolic activity of the tissues was monitored daily by the measurement of glucose concentration in the media supernatants of the three pools. The Glucose Liquid Reagent (Stanbio) was used for this, according to the manufacturer’s protocol with minor modifications: An amount of 57 μL of reagent was used and 3 μL of sample was added. Samples were incubated in 384-well microtiter plates for 10 min on an orbital shaker. Absorbance readings at 520 nm were performed in a microplate reader (BMG Labtech Clariostar). Chip culture medium and glucose feeding solution were measured daily as controls. The measured glucose levels of the glucose feeding solution were used as a correction factor to correct for evaporation effects of the culture medium during storage.

## Analysis

### RNA isolation and qPCR

Tissue equivalents were collected for RNA isolation using the NucleoSpin RNA kit (Macherey-Nagel GmbH & Co. KG). The cDNA was synthesized by reverse transcription of 150 ng total RNA (TaqMan^®^ Reverse Transcription Reagents, Thermo Fisher Scientific). The qPCR experiments were conducted using the QuantStudio 5 Real-Time PCR System 384-well Block and the SensiFAST SYBR Lo-ROX Kit, Bioline, according to the manufacturer’s instructions. The qPCR primers are shown in Table S 1. Positive control cDNAs for the human total adult small intestine, liver, brain (Zyagen) and kidney (BioChain) were used as comparison.

Statistical analysis of the gene expression data was performed with Prism 7.03 software (GraphPad) by presuming a log-normal distribution. A one-way ANOVA with Tukey’s multiple comparisons test was performed to compare the relative gene expression between different time points of the chip culture for each organ equivalent. P values are given at a 95 % confidence interval and considered significant < 0.05 (* = < 0.05, ** = < 0.01, *** = < 0.001). The geometric mean with geometric standard derivation is plotted.

### Immunohistochemistry

Immunohistochemical analyses were performed by staining with the following primary antibodies: hepatocyte nuclear factor 4 alpha R&D, mouse anti-human), CDX2 (BioGenex, mouse anti-human), Cytokeratin 8/18 (Santa Cruz Biotechnology Inc., mouse anti-human), Ki-67 (eBioscience, mouse antihuman), TRA-1-60 (eBioscience, mouse anti-human), TUBB3 or beta-3 tubulin (eBioscience, mouse anti-human), Na^+^/K^+^-ATPase (abcam, rabbit anti-human) (abcam, rabbit anti-human), Nanog (R&D, goat anti-human), Oct-3/4 (R&D, goat anti-human), vimentin (Thermo Fisher Scientific, rabbit antihuman), SLC10A1/NTCP (abcam, rabbit anti-human), PAX6 (BioLegend, rabbit anti-human), aquaporin1 (abcam, mouse anti-human), MRP2 (Thermo Fisher Scientific, rabbit anti-human), ZO-1 (Proteintech, rabbit anti-human), albumin (sigma, mouse anti-human) and nestin (eBioscience, mouse anti-human). The Apo-Direct Apoptosis Detection Kit (Thermo Fisher Scientific) was used for the detection of apoptotic cells, according to the manufacturer’s instructions.

Briefly, representative central 7 μm cryosections of the tissues were fixed in acetone at −20 °C for 10 min and washed twice with PBS. Tissue sections were then incubated with the respective primary antibody in 10 % goat or donkey serum in PBS for 2 h and washed afterwards twice with PBS. CF488A goat anti-rabbit IgG, CF594 goat anti-mouse, CF488A donkey anti-mouse and CF594 donkey anti-goat (all purchased from Biotium) and DAPI (Sigma-Aldrich) were used for visualization. Images were obtained using a Keyence BZ-X700E fluorescent microscope.

### RNA sequencing

A total of 16 RNA samples were analyzed with the NextSeq 500 sequencer (Illumina) to study RNA expression. The Illumina TruSeq stranded total RNA protocol was used including rRNA depletion (Cat. No. RS-122-2201). Cluster generation and sequencing was carried out by using the Illumina NextSeq 500 system with a read length of 75 nucleotides, according to the manufacturer’s guidelines. Sequence reads that passed the Illumina quality filtering were considered for alignment. Raw FASTQ files were checked for quality, aligned with the STAR mapper [Reference PMID 23104886] (2.5.3a) to the human genome assembly version GRCh37 and transcripts were quantified using Salmon [Reference PMID 28263959] (0.9.1). The resulting quast files were subjected to DESeq2 [Reference PMID 25516281] (1.18.1) analysis using the R statistical language. Samples were normalized and the dispersions were estimated using the default DESeq2 settings. DESeq2 was then used to identify differentially expressed genes for each pairwise comparison between organ and iPSCs. KeyGenes (www.keygenes.nl)^24^ was used to predict the identity of the ADME-MOC co-cultivated iPSC-derived samples.

## Results

### ADME-MOC design

An ADME-MOC design was realized with constraints of the scaled-down human physiology. The chip was laid out to host cell culture spaces for intestinal organoids, liver equivalents, renal organoids and neurospheres in an arrangement mimicking the *in vivo* situation (Figure 1 A). This combination of organs was chosen to allow for studies evaluating compound ADME profiles. Neuronal tissue was included as a fourth effector organ. Dimensional data of channels, cell culture compartments and flow characteristics were modelled to mirror the physiological relationship between them. In combination with the chip-pharmacokinetics model prepared, this enables not only the evaluation of the ADME profile of a compound, but also allows an extrapolation of the results to humans.

It was shown previously that renal cell differentiation and function was enhanced by the application of shear stress^25^. A separate circuit for renal cell culture was established to allow for adjustable shear rates on both sides of the kidney’s tubule. Hence, the layout comprised two circuits (termed “surrogate blood circuit” and “excretory circuit”) which overlapped in the kidney compartments and were separated by a porous cell culture-treated polycarbonate membrane (Figure 1 A). One medium reservoir compartment in each circuit allowed sampling of supernatants. The medium was perfused through the microfluidic network by two incorporated, pneumatic micropumps – one for each circuit. The pumping technology was based on TissUse’ original MOC platform^14^. Long-term stability and adjustability to various flow rates had been validated previously^11 12, 26^.

A parallel, physiological-inspired flow scheme through the compartments and a medium flow partitioning mimicking physiological ratios of blood flow were adapted for further quantitative *in vitro* to *in vivo* extrapolation (QIVIVE) (Figure 1 B). Therefore, the dimensions of the channels of the blood circuit were based on scaled human physiology. Data for this were obtained from 1,000 simulated male Europeans aged 40 to 60 and was gratefully provided by Certara/SimCyp. Average values for the four target organs are given in Table 1. The target organs received a distinct percentage of the blood flow from the main channel derived from the physiological data (Figure 1 C). This ratio was adjusted by the channel width and length, which both contribute to the channel’s flow resistance. Modelling of the resistances was carried out with MATLAB (Mathworks) and COMSOL Multiphysics 5.2a. Consequently, the flow was simplified to be laminar and steady. Hydrostatic pressures and any interaction of the two circuits were neglected.

**Table 1:** Parameters of key organs averaged from simulated data of 1,000 male Europeans. The ratios were calculated disregarding the organs not modeled in the chip. Only bold figures are totaled. The blood flow through the hepatic portal vein was simplified to be equal to the blood flow through the small intestine.

| Organ | Blood flow [L/h] | Ratio of blood flow [%] |
| --- | --- | --- |
| Small intestine | **64.3** | **35.3** |
| Liver | 82.0 | 45.2 |
| through portal vein | ~64.3 | 35.3 |
| through hepatic artery | **~17.7** | **9.9** |
| Kidney | **61.1** | **33.7** |
| Brain | **38.6** | **21.1** |
| **Total (arterial blood)** | **181.7** | **100** |

The compartments for the intestinal and neuronal equivalents were adjusted to fit standard 24- and 96-well cell culture inserts, respectively. The compartment for the liver equivalent was chosen to have a volume of 14.7 μL (3 mm filled height). This is in accordance with the downscaling of the other organs.

Both circuits contained two functionally separated compartments at overlapping positions: One for the cultivation of podocytes – to emulate the renal glomeruli – and one for the cultivation of epithelial cells of the renal proximal tubule. This segregation of the kidney’s functional units was adapted to account for substance filtration and reabsorption into the blood circuit. In contrast to the liver, the volume or number of cells does not define the renal functional output but rather the surface area that they provide for filtration and reabsorption. The complete surface of the human proximal tubule of a pair of kidneys is estimated to be 40 to 80 m^2^ when considering the epithelial cell’s microvilli surface^27^. The microvilli increase the surface by a factor of 30 to 60. Scaling the kidney in accordance to the ten liver lobuli, the surface of the chip’s tubules compartment had to remain between 6.7 and 26.7 mm^2^. We chose a surface of 13 mm^2^ considering the general size constraints of the chip. A single human glomerulus has a filtration surface of 0.17 to 0.21 mm^2^^28, 29^. Both human kidneys contain together approximately 1.5 to 2 million glomeruli^30^. By applying the same scaling factor, the glomerular compartment should have a surface of 2.6 to 4.2 mm^2^ in the MOC. We chose an area of 4 mm^2^ for the chip design. The output of the chip’s micropump was adjusted to create an average shear stress of 0. 1 Pa (1 dyn/cm^2^) on the surface of the epithelial cells of the proximal tubules (Figure 1 D)^31^.

### Human iPSC differentiations

All differentiated tissues used for ADME-MOC co-culture were derived from the same iPSC line, StemUse101, to ensure an autologous culture of different tissues (Figure 2). Liver, intestinal and neuronal tissue models were established and characterized as single static cultures before insertion into the ADME-MOC (Figure S 5– S 7). Similarly, renal organoids were produced in static culture, but cells were dissociated on day 12 of differentiation and transferred into the renal circuit of each ADME-MOC six days prior to the start of the co-culture experiment (Figure 3 A). Subsequently, renal organoids formed in the glomerulus kidney compartment. However, the renal cells did not attach in the tubules compartment of the kidney. The application of compatible coatings will be necessary to improve the attachment of renal cells. Further studies will address this issue. This was clearly not ideal, but we tolerated the cell loss. However, the renal organoid in the glomerulus compartment was believed to suffice for demonstrating the principal functionality of the differentiation strategy.

**Figure 2:**
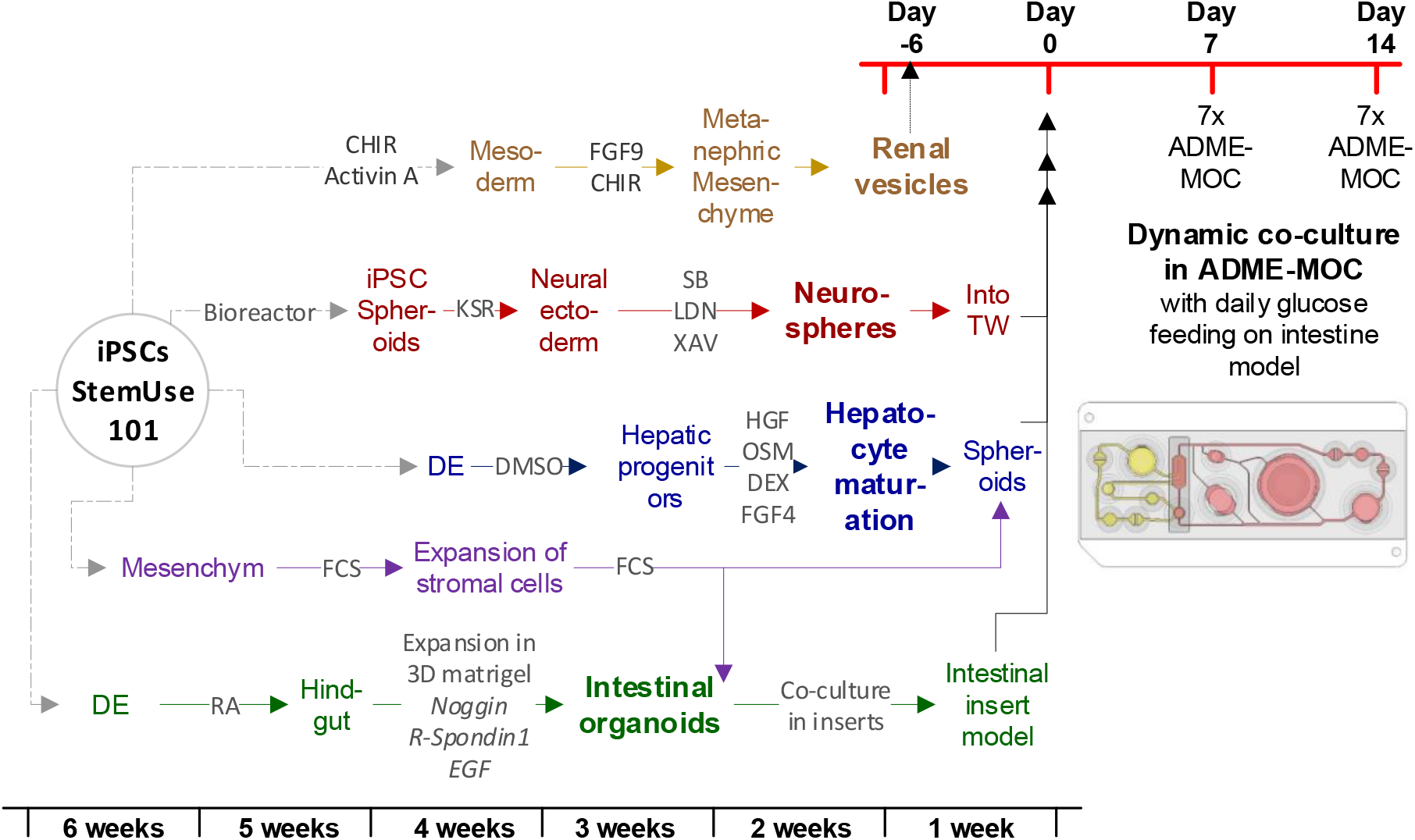
Schematic time frame of the iPSC differentiation into kidney organoids, liver spheroids, neurospheres and intestinal organoids, transfer and co-cultivation in ADME-MOC. DE: definitive endoderm.

**Figure 3:**
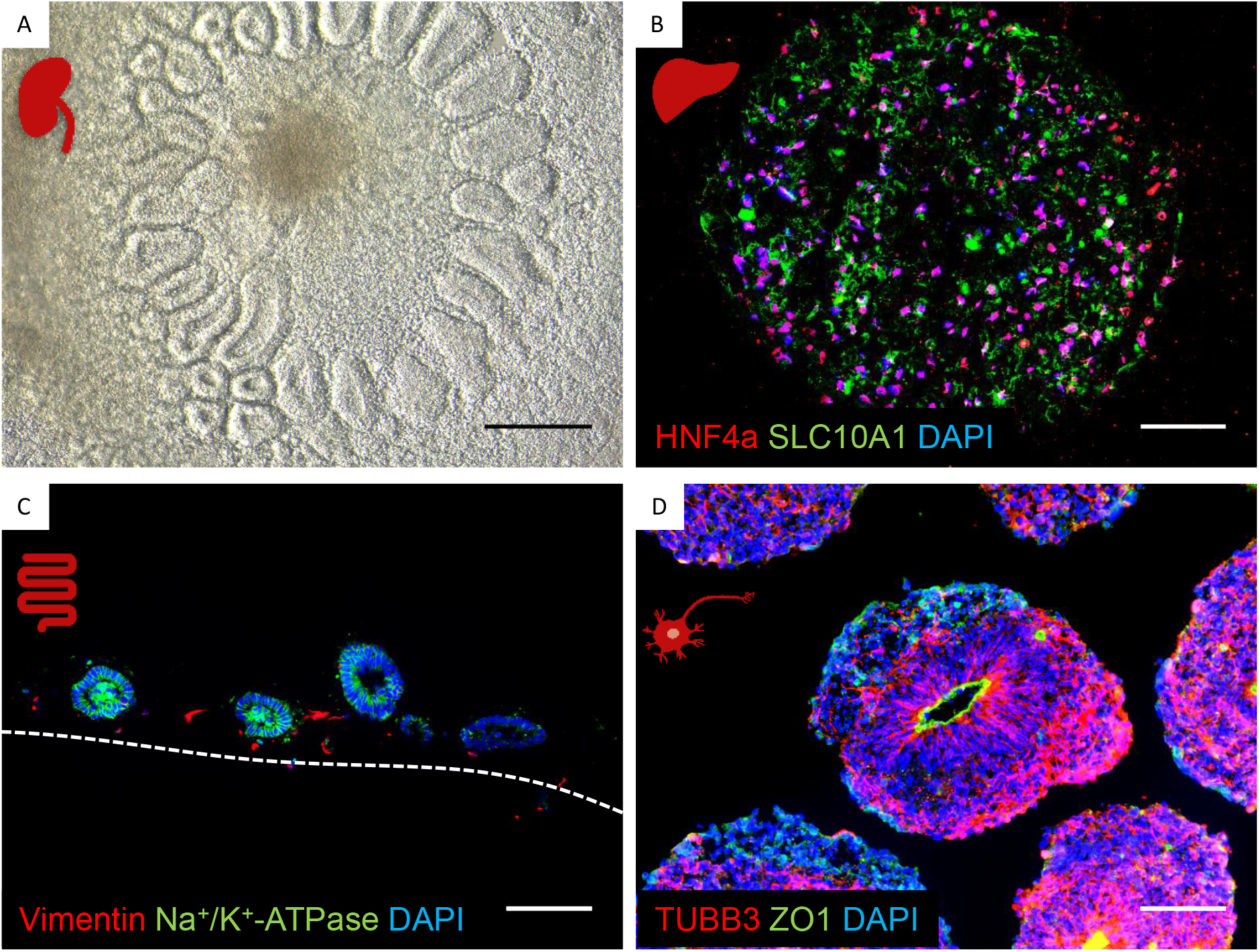
iPSC organoids on day zero of ADME-MOC culture A: kidney organoids before transfer into the ADME-MOC (scale 250 μm); B: transcription factor hepatocyte nuclear factor 4 alpha (red) and liver bile acid transporter SLC10A1 (green) expression in liver spheroids (scale 100 μm). C: Na^+^/K^+^-ATPase transporter (green) and stroma cells (vimentin – red) expression in intestinal insert model (scale 100 μm); D: neuronal spheroids stained for TUBB3 (red) and tight junction marker ZO-1 (green) (scale 100 μm).

It was shown previously^32, 33^ that stromal cells support hepatocyte function over prolonged culture periods. Therefore, liver equivalents were established by combining iPSC-derived hepatocytes with iPSC-derived stromal cells in a 3D spheroid model. Hepatocyte nuclear factor 4 alpha, a transcription factor which regulates the expression of several hepatic genes, and SLC10A1, a liver bile acid transporter, were observed by immunohistological staining in liver equivalents (Figure 3 B). Moreover, a co-culture of hepatocytes and stroma cells led to further differentiation, as indicated by significant upregulation of albumin and MRP2 (Figure S 6 A). Interestingly, a bile-salt export pump was only expressed in the spheroids and not detected in hepatocytes in monolayer (Figure S 6 A).

We used standing 24-well cell culture inserts for the intestinal barrier model to establish a permeable barrier separating the apical feeding space and the blood perfusion circuit in the ADME-MOC. An iPSC-derived stromal cell bed was combined here with preformed intestinal organoids. Stromal cells were stained with vimentin and intestinal organoids with Na^+^/K^+^-ATPase (Figure 3 C). The intestinal organoids retained their spherical shape with characteristic marker expression, whereas stromal cells distributed equally around the tissues.

Neurosphere differentiation was performed in a bioreactor system (DasBox) over 32 days. The neurospheres showed neural tube-like structures with a distinct ZO-1 positive lumen surrounded by positive TUBB3 cells (Figure 3 D). Additional results for iPSC and differentiated tissue characterization can be found in the Supplement Figure S 5-7.

### Establishing the ADME-MOC co-culture

All tissue models were established in their specific iPSC culture media prior to their insertion into the ADME-MOC. However, a common circulating medium supporting all tissues was mandatory to allow for cross talk between organoids via cytokines and other molecular communication modes. Therefore, a common, growth factor-depleted medium was used in the surrogate blood circuit during co-culture. The excretory circuit contained a similar medium which was additionally devoid of human AB serum and ITS. Neither media contained glucose, which was supplied exclusively to the apical side of the intestinal tissue model. All four organ models were sufficiently supplied with glucose due to the absorption through the intestinal model and subsequent circulation of the medium in both circuits (Figure 4 B).

**Figure 4:**
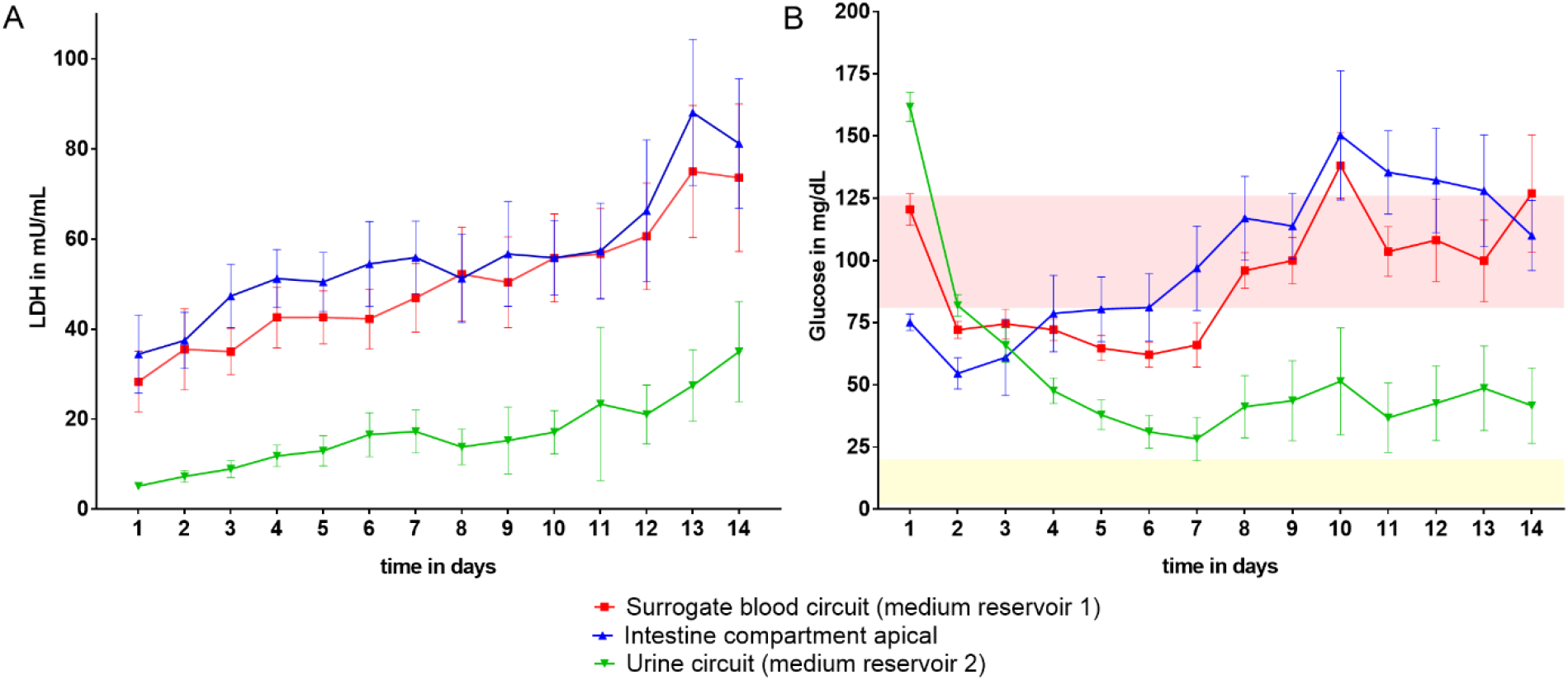
Systemic tissue viability in the ADME-MOC over 14 days of co-culture measured by LDH activity (A) and glucose balance (B). The physiological glucose concentration range in the human blood^48^ (red area) and urine^49^ (light yellow area) of healthy people is drawn for comparison. The means with 95 % confidence interval are plotted. N = 28 until day 7 and N= 21 until day 14.

The medium was perfused through the microfluidic network by two integrated pneumatic micropumps – one for each circuit – providing physiological pulsatile fluid flow at a microliter scale. The main flow rate in the blood circuit was 16.9 ± 0.7 μL/min. This led to an approximate medium turnover time of 1. 35 h. The excretory circuit had a flow rate of 6.6 ± 0.3 μL/min and a turnover time of 1.45 h. The minute amounts of enriched cultivation medium within the two circuits enabled cross talk between the tissues.

The μPIV analysis revealed correct perfusion of the intestinal and neuronal equivalents according to the criteria desired. Interestingly, the liver and the kidney equivalents were over- and undersupplied, respectively. The reasons for this phenomenon could evolve around the fact that the two circuits are in exchange with another. Additionally, the various inserts should be considered more appropriately in the modeling.

### Establishing the iPSC-derived organoid co-culture in the ADME-MOC

Co-culture viability and performance was assessed daily by analyzing metabolic products in the culture medium (Figure 4) and microscopic inspection. This was feasible through the chip’s transparent bottom allowing live tissue imaging.

Overall cell viability was measured as total LDH release into the media (Figure 4 A). Therefore, samples were taken daily from the surrogate blood circuit, the excretory circuit and the apical intestinal compartment. Due to the high cell turnover in the intestinal organoids and the kidney organoids, we saw an increasing release of LDH into these medium supernatants (Figure 4 A). However, compared to the positive control (1,013 mU/mL) where all cells in the ADME-MOC were lysed and LDH was measured, the LDH release during the 14 days of co-culture was low, reflecting physiological cell turnover in the chip. Staining for TUNEL/Ki67 showed a high viability in the liver, intestinal and renal model on day 14 of co-culture; only the neuronal model showed some dead cells (Figure 5 C, F, I, L).

**Figure 5:**
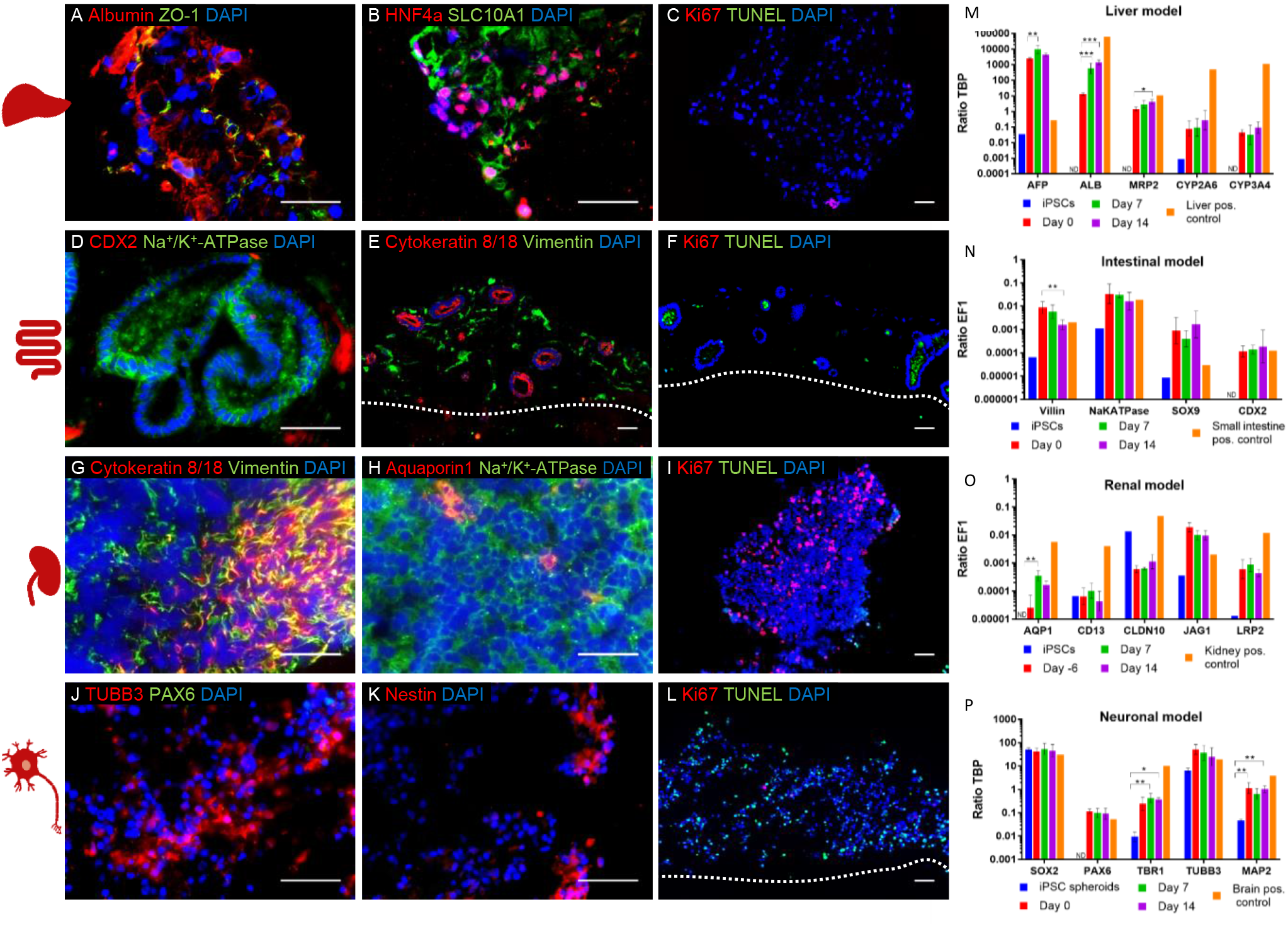
Characterization of the liver, intestinal, renal and neuronal model co-cultivated in the ADME-MOC for 14 days. Immunostaining A – L: A - C liver model: A: albumin and ZO-1, B: Hepatocyte nuclear factor 4 alpha and SLC10A1, C: Ki67 and TUNEL. D – F: intestinal model: D: CDX2 and Na^+^/K^+^-ATPase, E: cytokeratin 8/18 and vimentin, F: Ki67 and TUNEL. G – I: renal model: G: cytokeratin 8/18 and vimentin, H: aquaporin 1 and Na^+^/K^+^-ATPase, I: Ki67 and TUNEL. J – L: neuronal model; J: TUBB3 and PAX6, K: nestin, L: Ki67 and TUNEL. Scale 50 μm. M –P: qPCR results of the co-culture of the iPSC-derived intestinal, liver, renal and brain model in the ADME-MOC. Data are normalized to the housekeeper EF1 or TBP. N = 3 – 6 for day 0, 7 and 14, N = 1 – 3 for undifferentiated iPSCs and N = 1 for the positive controls represented. ND = no signal detected. Ordinary one-way ANOVA with Tukey’s multiple comparisons test was used for statistical analysis. (95 % confidence interval, * = < 0.05, ** = < 0.01, *** = < 0.001.) The geometric mean with geometric standard derivation is shown.

Similarly, glucose concentrations were measured to determine whether all tissues were supplied sufficiently. Feeding glucose only apically through the intestinal equivalent was shown to be sufficient to supply both the surrogated blood circuit and excretory circuit over 14 days of chip co-culture (Figure 4 B). Additionally, the glucose values of the surrogated blood circuit and excretory circuit were in the range of physiological glucose concentration of the human blood and urine, respectively.

All organ equivalents were assessed on the protein and gene expression level on day 14 of co-culture to determine their identity, functionality and viability (Figure 5).

The liver equivalents showed a high expression of albumin. The tight junction protein ZO-1 was also expressed in the cell membranes (Figure 5 A). The hepatic transcription factor HNF4a was detected in the nuclei and SLC10A1, a liver bile acid transporter, was visible (Figure 5 B), conjointly illustrating the hepatic character of the liver equivalent. The expressional data of the liver equivalents showed a significant increase of albumin and MRP2 from day 0 until day 14 of the ADME-MOC co-culture (Figure 5 M). By contrast, no liver-specific genes were detected in undifferentiated iPSCs (Figure 5 M). The mRNA expression of albumin, CYP2A6 and CYP3A4 were much lower in the ADME-MOC compared to the human adult positive control. Therefore, we assumed that the liver equivalents were not completely matured. This was also indicated by the expression of APF, the fetal form of serum albumin. It was expressed more highly in the ADME-MOC liver model than in the liver adult positive control. By contrast, MRP2 (multidrug resistance protein 2), an important transporter on the apical membrane of bile canaliculi in hepatocytes, had a comparable mRNA expression to the liver adult positive control. RNA sequencing results revealed an upregulation of many cytochrome P450 genes in the iPSC-derived liver model in comparison to the iPSCs (Figure S 10 and Figure S 11). Interestingly, only CYP1A2 was not upregulated from iPSCs towards the liver model.

Intestinal organoids have a lumen and express the Na^+^/K^+^-ATPase transporter at high rates. Few cells within the organoids were positive for the intestinal transcription factor CDX2 (Figure 5 D). Cytokeratin 8/18 was only expressed in the epithelial cells of the organoids and vimentin only in mesenchymal cells in the stromal bed around the organoids (Figure 5 E). Stable steady-state gene expression of Na^+^/K^+^-ATPase, SOX9 and CDX2 (both intestinal transcription factors) over 14 days in the ADME-MOC was also shown by qPCR analysis in the intestinal model. Villin and Na^+^/K^+^-ATPase gene expression on day 14 were comparable to the positive control of the human adult small intestine, although it was downregulated from day 0 to 14. The intestinal transcription factors SOX9 and CDX2 were expressed more highly in the co-cultures from day 0 to 14, just as in the small intestine positive control. This indicated that the cells were still in the maturation process. As expected, all markers had a much lower expression in undifferentiated iPSCs (Figure 5 N). RNA sequencing results revealed an upregulation of many genes involved in the Wnt signaling pathway, which is important for intestinal development, in the iPSC-derived intestinal model in comparison to the iPSCs (Figure S 13).

As expected, kidney organoids showed a mixed culture of epithelial cells (cytokeratin 8/18) and mesenchymal cells (vimentin) (Figure 5 G and H); the transporter Na^+^/K^+^-ATPase was highly expressed selectively in the cell membranes (Figure 5 H) and the proximal tubules marker aquaporin 1 was only expressed in a few cells (Figure 5 H). Additionally, we saw a high number of proliferating cells in the kidney organoids and only a few apoptotic cells (Figure 5 I). Gene expression data showed a clear progression of the differentiation without achieving maturation (Figure 5 O). Aquaporin1 (a water channel), for example, is significantly upregulated from day -6 to day 7 but is still expressed lower than in the adult kidney positive control. The alanine aminopeptidase (CD13), an enzyme in proximal tubular epithelial cells, was expressed but also lower than in the adult kidney positive control. Claudin 10, a tight junction component in the renal tubule and glomerulus, was surprisingly highly expressed in undifferentiated iPSCs. This might have occured due to iPSCs growing in colonies with high cell-cell contact including tight junctions. Jagged1 (Jag 1) is involved in early nephron development; therefore, we saw a higher expression in kidney organoids on day -6 with a tendency to downregulated until day 7 and 14. The lower expression of the kidney positive control indicates that the differentiation status of kidney organoids was not mature. Megalin, also known as low-density lipoprotein-related protein 2 (LRP2), is a receptor in many absorptive epithelial cells and exhibits a static expression over the course of 14 days. However, there was a high difference to its equivalent in the adult positive kidney control.

The neurospheres self-assembled inside the Transwell systems into larger structures during the 14 days of ADME-MOC cultivation. Tissues appeared to be heterogeneous with distinct TUBB3 and nestin positive areas (Figure 5 J and K), which indicates structural changes of the brain equivalent induced by the ADME-MOC culture. No PAX6 positive cells were observed after 14 days, indicating a decline of the neural stem cell fraction. Brain equivalent RNA expression showed a stable expression of the neural stem cell markers SOX2 and PAX6, the neuronal markers TUBB3 and MAP2, and the cortical marker TBR1 from day 0 to day 14 of the ADME-MOC culture (Figure 5 P). Neural stem cell and neuronal markers were expressed at a comparable level to primary brain tissue, while TBR1 was expressed lower. This implies that the cortical developmental process was not completed when the ADME-MOC co-culture was started. RNA sequencing results revealed an upregulation of many genes important for axon guidance and neuron development in the iPSC-derived neuronal model in comparison to the iPSCs (Figure S 12).

Principle component analysis (Figure 6) and KeyGenes prediction from RNA sequencing data (Figure S 9) showed that the models not only maintained their phenotype in the common media, but also even differentiated further along their destined paths.

**Figure 6:**
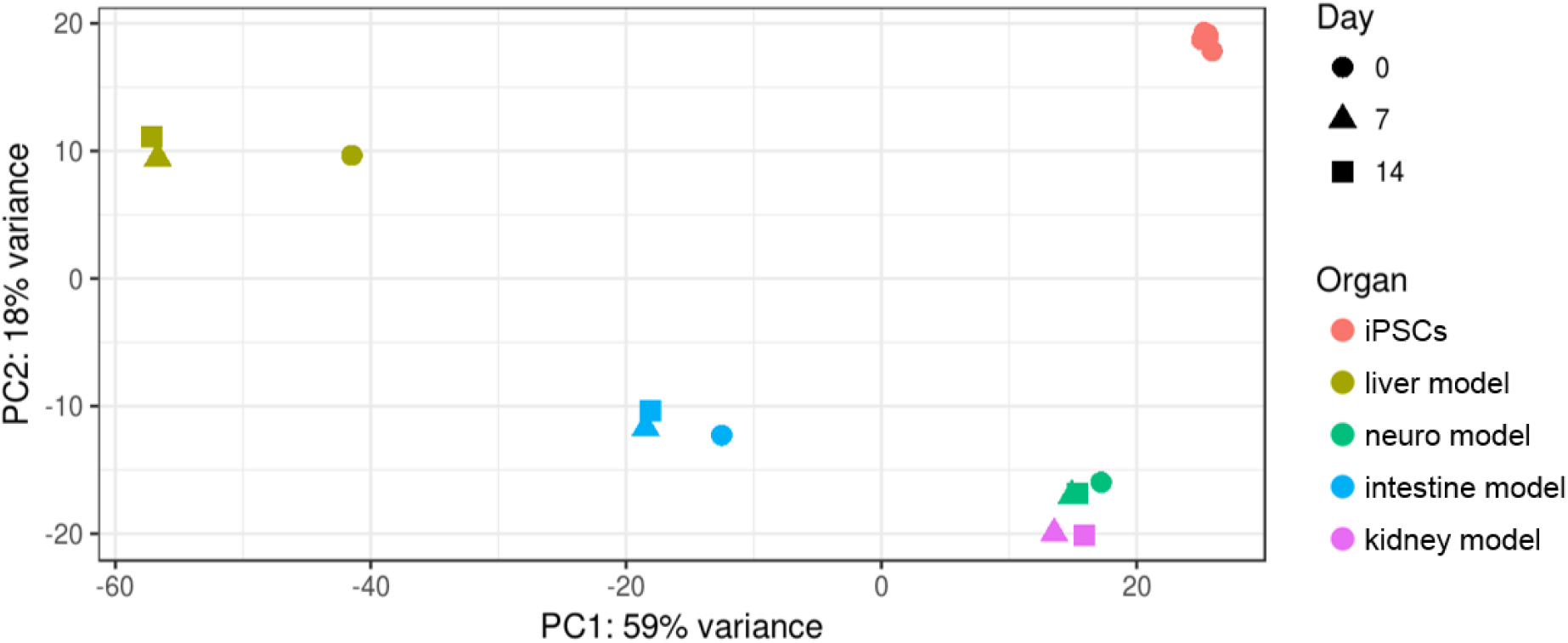
Principle component analysis on RNA sequencing data of the liver, intestine, brain and kidney model cultivated over 0, 7 and 14 days in the ADME-MOC. Five different human iPSC lines were analyzed for comparison.

RNA sequencing and the following principle component analysis on all four different iPSC-derived models co-cultivated over 14-days in the ADME-MOC, and five different iPSC lines were analyzed regarding the first two principal components (Figure 6). The first component explained most of the variance. Distinct clustering of each model was visible. Interestingly, the kidney and brain model clustered close to each other. The shift from day zero to day seven and fourteen was striking for the liver and intestine model. Co-culture in the ADME-MOC under dynamic conditions was, therefore, having a profound effect on the RNA expression profile. This was also confirmed by the high throughput barrier multiplex qPCR analysis of the intestinal and kidney models (Figure S 8). Investigation of 94 different barrier genes showed 49 significantly different regulated genes (Figure S8 A). Distinct clustering of the intestinal versus the renal models was also obvious in the principal component analysis (PCA) (Figure S 8 B).

The RNA sequencing data from the ADME-MOC experiments were used for comparative analysis using KeyGenes. It is an algorithm to predict the identity and to determine the identity scores of queried samples to provided transcriptional profiles of different organs or cell types^34, 35^. The algorithm uses the top 500 most differentially expressed genes of the available data sets and matches them with the samples provided to predict a tissue type. Different human transcriptional signature sets are available – fetal, 1st trimester fetal, 2nd trimester fetal and adult.

The intestine, liver and brain model were identified by KeyGenes using the human adult training set (Figure S 9 A and B). The intestine model co-cultivated over 14 days in the ADME-MOC showed the highest identity score with the intestine when using the adult training set (Figure S 9 A). Only the kidney model was not predicted appropriately; instead, it was predicted as a brain sample. This is probably due to an incomplete renal differentiation and a coexisting differentiation into neuronal cells in addition to the occasional renal organoids.

The liver model showed a higher identity score when the human fetal training set was used instead of the human adult training set (Figure S 9 B), which is also in accordance with the gene expression data. Therefore, we assume that the liver model has a fetal phenotype. Moreover, the neuronal model also had a higher identity score in the fetal training set. RNA sequencing data for all four organ models were additionally evaluated by gene ontology (GO) analysis. Significant terms for biological processes, molecular function and cellular compartment can be found in the Supplement Table 2.

## Discussion and conclusion

### Scaling

There are different approaches for downscaling proceedings in MPS. Researchers used to biological contexts are probably most familiar with allometric principles that explain the evolutionary scaling relationships between small and big animals. While allometric scaling can be helpful in animal to human extrapolation^36, 37^, modeling the human physiology in a miniature format is scaled differently^38^.

Ultimately, the emulated human physiology should perform according to its original counterpart. Thus, scaling should be met by function and not by mass^39, 40^. If the *in vitro* liver produces less albumin than the same volume of liver in a human body, for instance, the volume of the *in vitro* equivalent should be increased. This bears two obstacles. Firstly, organs such as the liver perform several functions that are not proportionally under- or over-realized in *in vitro* models – albumin might be low, while cytochrome activity might be average. This can only be met by compromises and the focus on a subset or the physical separation of the original functions that ought to be scaled. Furthermore, the function must be assessable and attributable to the scaled organ – especially in a system with several cell or organ types.

Secondly, when creating a multi-organ MPS, there needs to be an agreement on a single unit within the chip that acts as a scaling guideline for the rest of the chip. Within our group, we normally use the equivalent of ten liver lobules as the basis for the scaling. As laid out in other works, each organ can be subdivided into functional organoids^41^. The liver contains approximately one million liver lobules – each performing the entirety of the liver’s capabilities on a microscale. Keeping the number of organoids in a multi-organ MPS at similar proportions, one would continue to incorporate the equivalents of five pancreatic islets, 3,000 lung alveoli and so on and so forth. The aim is to model a biological systemic interplay of human organ equivalents in a small format. Again, if we are technologically unable to recreate the functional units, we must match the cell number or surface area to the functionality required. Of course, the ratios in which the organoid numbers are defined might have to be adapted when it comes to modeling different ages or certain pathologic conditions. Liver and kidney, for example, usually scale differently in infants than in adults.

We also paid attention to the flow rates within both circuits in the current design of the ADME-MOC. We have been unable to scale the blood’s volume with the same factor as the individual organ equivalents, as it would lead to volumes of some 60 to 80 μL. In fact, we would have to match the volume to the solubility of the medium to match the blood’s capability of transporting oxygen. Instead, we scaled the flow rate to match the *in vivo* demand and kept the ratios with which the individual organs were supplied. Thus, each organ equivalent was exposed to flows and concentration conditions derived from its organoid counterpart.

Admittedly, the scaling applied cannot recapitulate the physiology perfectly due to technical, biological and physical limitations. In fact, we would argue that there is no sound scaling enduring all critique. If an ‘organismic’ arrangement of organ models is self-sufficient and homeostatic, the fine-tuning and correct scaling will be adopted by the models themselves. The liver, for example, should inherently be able to adapt its activity (and volume) to the demands of the other organ models. Considering the shortcoming of a generally applicable scaling, coherent organoid numbers, flows of liquids and gradients of substances are the real constrains of these systems. Admissible parameters, read out for pharmacology and toxicology, will, eventually, be extrapolated from the chip’s organoid level to human size by physiologically based pharmacokinetic modeling rather than allometry. The groundwork for this has been laid out here.

### Chip cultivation

To the best of our knowledge, this study is the first successful long-term co-cultivation of four different iPSC-derived tissues from a single donor cultured in a physiologically based pharmacokinetic compliant MPS. Those tissues show defined marker expression towards the directed tissue. Furthermore, we would like to point out that no growth factors were added to the medium during the co-culture in the ADME-MOC. The basic medium with 5 % human AB serum was sufficient for all four human iPSC-derived tissues. There was no dedifferentiation observed; we could even remark an advanced maturation of those different tissues over time.

Nevertheless, a few obstacles remain: The tissue maturation is not fully attained and further developments considering medium composition, scaffold and co-culture with stabilizing and supporting cells are necessary before the start of the future chip co-cultures. The state-of-the-art iPSC differentiation protocols performed mostly lack the generation of fully matured organ models and rather produce fetal tissue models.

Furthermore, we aspire towards a closed, iPSC-derived endothelialization of the channels and tissues in the chips. This would support the use of full blood for nutrition and oxygen supply.

Vunjak-Novakovic and colleagues are currently working on the so-called HeLiVa platform to combine iPSC-based vascular, liver and cardiac microtissues^42^. To the best of our knowledge, the combination of all three tissues has not yet been published, although they show great maturation of iPSC-derived hepatocytes and cardiomyocytes by small molecule screening^43^ and electromechanical conditioning^42^, respectively.

### Tissue performance

We could show that the combination of stromal cells with the hepatocytes support the maturation of liver equivalents revealed by the significant upregulation of albumin and MRP2 (Figure S 6). This maturation of hepatoblasts as a result of the co-culture with fibroblasts was also shown by Shan and colleagues, who used a mouse feeder monolayer^43^. Additionally, the co-culture with the intestinal, renal and neuronal models in the ADME-MOC led to further maturation of the liver model (Figure 5). However, the high expression of Alpha-Fetoprotein and the lower expression of CYP2A6 and CYP3A4 in comparison to the adult liver control shows that there is no full maturation of hepatocytes but rather a fetal phenotype and function of the liver equivalent. KeyGenes prediction also confirms the assumption that the iPSC-derived liver model shows more similarities with a fetal than an adult liver (Figure S 9 A and B). Additionally, the comparison of metabolization enzymes to adult liver RNA is challenging due to high donor variability and individual stages of induction. We are confident that the development of improved differentiation protocols regarding medium, growth factors, metabolism and scaffolds will lead to more mature cells and tissues.

Regarding the intestinal model, the highest consent of the KeyGenes prediction can be seen between the 14-day intestinal co-cultures model and the adult intestine (Figure S 9 A). This reflects the previous observation that the co-culture under dynamic conditions in the ADME-MOC has a maturation effect on the intestinal model. We seek to obtain an intestinal epithelial cell layer that is opened at the luminal side, apically facing the top of the Transwell, while the basolateral side lies on the membrane of the Transwell to further improve our intestinal model. A similar single organ approach has recently been shown with iPSC-derived and primary small intestinal organoids in a chip system^44, 45^. Moreover, a dense tight layer of endothelial cells underneath the Transwell should seal the barrier to the blood surrogate.

The kidney model was the only tissue that was not successfully predicted by KeyGenes. Instead, KeyGenes predicted a brain or muscle organ from the adult of fetal training set, respectively (Figure S9 A and B). We assume that the rare renal organoids, which arouse during differentiation, were overgrown by coexisting cells. Therefore, the differentiation efficiency into purer renal organoids is to be optimized. Additionally, we want to use more defined and specialized cells for the different parts of the kidney compartments in future studies. The glomerulus is proposed to house filtering renal cells, such as podocytes, on one side and endothelial cells on the other. Furthermore, the tubular loop should receive proximal tubular epithelial cells, which reabsorb solutes and water back into the blood circuit, which is also ideally lined by endothelial cells.

Although the chip design and the organoids on top of the glomerulus support the tightness of the porous polycarbonate membrane, the biological barrier function of the intestine equivalent and the renal equivalent was incomplete in this study, as there was no tight layer of endothelial or epithelia cells holding the barrier. Nevertheless, the technical barrier between the excretory circuit and the blood surrogate circuit could be maintained, as was shown by stable concentrations between the different media compositions. The excretory circuit, for example, contained much less LDH than the surrogated blood circuit (Figure 4).

Immunofluorescent staining showed an extensive amount of vital neuronal cells after 14 days of ADME-MOC co-culture. However, a considerable number of apoptotic cells was also observed (Figure 5 L). Daily media exchanges with glucose-free maintenance medium to remove waste products or the culture under air-liquid interface (shown by Tieng and colleagues^46^) to increase the oxygen supply of the tissue might improve the culture conditions of the neuronal model. KeyGenes prediction showed the close similarity of the initial neuronal culture to fetal tissue (Figure S 9 B), however, there is a decrease over time, which supports the assumption that culture conditions are not yet ideal. Once culture conditions have been improved, more mature neurospheres which contain neuroglia could be included to further increase the complexity of the model. Another significant step is the introduction of iPSC-derived blood-brain barrier-specific endothelial cells^47^ to the system, to truly separate the brain equivalent from the blood circuit and, thus, emulate the blood-brain barrier.

We believe strongly that the combination of tissue engineered autologous 3D heterogeneous organ models and the communication between those different tissues obtained through the connection of the organ models by the dynamic MPS circulation will lead to a significant improvement in the prediction of drug response in clinical studies. In addition, iPSC-derived tissue standardization will neglect the donor variabilities of primary tissues. The generation of high numbers of the standardized iPSC-derived tissues will eliminate the bottleneck of primary cells with limited tissue sources such as the liver or pancreas.

The assembly of tissues from one single population of iPSCs will ultimately enable the creation of a donor-on-a-chip for drug screening. Receiving tissues from a single donor will be even more important once immune components are present in the chip co-cultures. Hence, rejection reactions as a result of the systemic connection of tissues with different genetic background can be prevented. Furthermore, the generation of a patient-on-a-chip from a disease donor gives great potential regarding personalized medicine and allows, for instance, the focus on genetic mutations of a single donor. The patient-on-a-chip holds great promise for further investigations, especially in the emerging field of individualized cellular immunotherapies in oncology, autoimmunity and transplantation medicine.

## Author Contributions

1. Anja Ramme designed, performed, analyzed experiment and wrote manuscript
2. Leopold Koenig designed, performed, analyzed experiments and wrote manuscript
3. Tobias Hasenberg designed, performed experiments and wrote manuscript
4. Christine Schwenk performed experiments and qPCR
5. Corinna Magauer chip production
6. Daniel Faust performed qPCR
7. Alexandra K. Lorenz performed chip experiment
8. Anna Krebs performed chip experiment
9. Christopher Drewellperformed chip experiment
10. Kerstin Schirrmann performed flow calculation and chip design
11. Alexandra Vladetic performed multiplex qPCR
12. Grace-Chiaen Lin performed multiplex qPCR
13. Stephan Pabinger bioinformatic analysis of RNA sequencing data
14. Winfried Neuhaus analyzed multiplex qPCR and RNA sequencing data
15. Frederic Bois PK/PD modelling
16. Roland Lauster designed experiments, supervised and reviewed manuscript
17. Uwe Marx formulated the strategy for this study and reviewed the manuscript
18. Eva Dehne formulated the strategy, designed experiments and chip, wrote manuscript

## Acknowledgment

We would also like to show our gratitude to Philip Saunders, Peter Mangel, Dr. Mir-Farzin Mashreghi, Katrin Lehmann, Frederik Heinrich, Ilka Maschmeyer, Phenocell SAS and Certara/SimCyp. This work was supported by Stiftung SET zur Förderung der Erforschung von Ersatz-und Ergänzungsmethoden zur Einschränkung von Tierversuchen, project 060 to Winfried Neuhaus.

